# Discovering Disease Genes in PPI Networks: A Bridge from Centrality to Communities

**DOI:** 10.1101/2023.09.08.556873

**Authors:** Mehwish Wahid Khan, Rauf Ahmed Shams Malick, Hocine Cherifi

**Affiliations:** School of Computing, National University of Computer and Emerging Sciences,Karachi 75030,Pakistan; ICB UMR 6303 CNRS University of Burgundy Dijon, France

**Author notes:** These authors contributed equally to this work.

## Abstract

Targeted therapies have become pivotal in modern clinical oncology, driven by a molecular-level understanding of cancer’s intricacies, its progression, and innovative research and technology. Personalized and targeted treatments hinge on identifying key genes, hub genes, or biomarkers. Protein-protein interaction (PPI) networks are instrumental in understanding the molecular basis of diseases. While existing literature has identified significant genes based on network centrality, investigations based on community-aware centrality have been notably absent. This omission matters because disease networks frequently display modular structures, necessitating a new perspective. This study bridges the gap between network centrality and community-based investigations. Indeed, in modular networks, node influence can be categorized into two types: local impact within its community, determined by intra-community connections, and global effect on nodes in other communities, established through inter-community links. This concept extends conventional centrality measures to networks with a community structure. Initially, we performed a comparative analysis of seven PPI networks related to cancer and noncancerous conditions. We explore the correlation between classical network centralities and their equivalents at the global (inter-community) and local (intra-community) levels. Notably, we consistently observed a high correlation between network degree and local degree centrality in all PPIs, leading us to select local degree centrality for further investigation. Pronounced modularity characterizes prostate and cervical disease networks. Consequently, we investigate these networks to identify key genes at the local community level and validate them by examining their expression levels. Variations in gene expression between cancerous and non-cancerous tissues bolster our findings. We identify a novel set of genes as potential key players in prostate and cervical cancer. Specifically, in cervical cancer, the top genes at the mesoscopic level include AKT1, CDK2, BRCA1, VEGFA, SRC, PSMD14, MRPL3, TP53, and NUP37. Meanwhile, the top genes identified in prostate cancer are FOS, TP53, UBA52, HLA-B, TSPO, and CD19. Although we focus on cancer data, our methodology’s versatility makes it applicable to other disease networks, opening avenues to identify key genes as potential drug targets.

## Introduction

Cancer is a multifaceted disease influenced by many genetic and phenotypic factors that contribute to developing and regulating specific cancer types. While some genes are shared among various cancer types, others are unique to particular tumors. Researchers have recognized the significance of biological networks in cancer studies, focusing on specific network types such as Protein-Protein Interaction (PPI) networks, metabolic networks, gene-regulatory networks, and drug-protein networks. The primary objective of studying these diverse networks in cancer research is to pinpoint the genes that play crucial roles in cancer development at the pathway level. Simultaneously, hundreds of drugs have been developed to target either overexpressed genes or mutated in cancer. Despite abundant drugs and high-throughput data encompassing gene expression profiles, methylation information, and mutation details, cancer prognosis remains a formidable challenge. To address this challenge and improve the selection of drugs for cancer treatment, researchers have devised intelligent methods that consider the specificity of the target genotype or phenotype. These methods aim to identify the most responsive drugs by analyzing gene expression profiles. This approach is promising for enhancing cancer prognosis and personalized treatment.

In drug discovery and cancer research, Machine Learning and Deep Learning methods have become indispensable tools for identifying the most promising drug candidates from vast cell line samples and many available drugs [1]. When it comes to targeting specific tumor types, analyzing expression profiles from hundreds of cell lines has proven invaluable in identifying irregularities in gene expressions shedding light on potential drug targets. However, this method primarily relies on individual gene expression profiles, potentially overlooking specific genes’ critical roles within intricate biological networks. Recognizing this limitation, researchers have turned to holistic approaches to enrich our understanding of tumors’ genetic and phenotypic dimensions. These comprehensive approaches encompass the study of significant mutations, methylation patterns, copy number variations (CNVs), gene expression levels, hub genes, and driver genes. Yet, within this mosaic of factors, the structural position of a gene within biological pathways remains a pivotal consideration. For instance, an under-expressed hub gene, functioning as a regulator, can profoundly influence the pathways it governs. It’s worth noting that identifying hub genes within the context of Protein-Protein Interaction (PPI) networks holds great promise. These networks, which reveal the complex web of protein interactions within a cell, offer a more complete understanding of how genes function in concert. However, despite their potential, current methods for identifying hub genes come with their implications. This study delves into the nuanced landscape of PPI networks, exploring their role in identifying critical nodes in cancer-related pathways. By comparing control and cancer PPI networks within the same tissue, we aim to unravel the structural differences that may hold the key to more effective cancer treatments.

Cancer research has advanced significantly, unveiling numerous genes associated with various cancer types. However, not all genes are universal; some are specific to particular tumors. Studying biological networks, including Protein-Protein Interaction (PPI) networks, has become a pivotal tool in comprehending cancer mechanisms [2]. Researchers strive to pinpoint essential genes that drive cancer development alongside hundreds of drugs designed to target these genes. Despite the wealth of data and highthroughput technologies, selecting the most effective treatments remains challenging [3]. Current methods often rely solely on individual gene expression profiles, potentially neglecting the roles of genes within intricate biological networks. Holistic approaches incorporate mutations, methylation, copy number variations (CNVs), and the structural positions of genes within pathways to provide a more comprehensive perspective. Hub genes, typically identified through network centrality measures, play pivotal roles in biological networks. Various methods have been proposed to identify critical nodes, such as statistical approaches [4], [5], hypergraph models [6], and node ranking [7]. Considering network modularity is crucial [8,9]. Highly modular PPI networks may be less resilient to removing critical nodes, while less modular networks with more interconnectedness tend to be more resilient. This aspect must be considered when identifying essential nodes. Despite the abundance of centrality measures, current methods often fail to consider network modularity when identifying critical nodes. Additionally, differences in network resilience and modularity between control and disease PPI networks, particularly in cancer studies, have yet to be explored [10], [11].

The present study investigates how the PPI networks of control differ from cancer PPIs of the same tissue. The comparison is performed at the meso level and compared the attributes of communities accordingly. The objective of understanding the modular composition of the PPI among control vs disease is to primarily understand the role of critical nodes. Secondly, in the presence of non-overlapping communities, it might be essential to identify community-level critical nodes along with the network-level critical nodes. Our analysis of multiple cancer and control PPI networks showed the prevalence of some highly modular PPI networks with non-overlapping node-based communities. Particularly in Colon and Prostate cancers, it is evident that communities have low overlaps and must represent distinguished biological functions. Furthermore, our analysis reports that the network-level critical node analysis in the above mentioned cancer types might not be effective compared to the community-level centrality analysis. Multiple communities have respective hubs or critical nodes while performing specific biological functions. The study may open new avenues for general network biology-based critical node analysis to identify biological functional pathway-based critical nodes rather than considering nodes at the network level.

The structure of our paper includes an overview of the data and experimental design in the next section, followed by the presentation of results, a discussion of our findings, and, finally, the conclusion.

## Materials and methods

### Data assimilation and Network Construction

In the Protein-Protein Interaction (PPI) network, the nodes are the proteins, and the edges represent the interactions among those proteins. The PPI networks used for analysis are constructed from GenBank and UniProt by [12]. The data set comprises PPI networks of normal and cancerous affected colon, prostate, breast, lung, ovarian, prostate, and cervical tissues. The PPI network pairs represent each cancer type in the present study.

### Experimental Description

We perform the network analysis on all fourteen cancerous and non-cancerous PPI networks. At the network level, we investigate multiple centralities referred to as classical measures. They include degree, closeness, betweenness, page rank, Katz, and subgraphs. The Infomap algorithm identifies communities within the PPI networks to compare the mesoscopic and macroscopic centralities. Infomap is a graph community detection algorithm known to achieve high-quality communities [13, 14]. It uses minimum description length (MDL) to find the quality of detected community results. Community-aware centrality measures include two components: a global and a local community-aware centrality measure [15, 16]. Given a classical centrality measure, one computes the local component, considering the communities as isolated networks. One forms the network of inter-community links with their attached nodes to compute the centrality global component. To evaluate the similarity between the corresponding classical, local, and global centralities, we compute the Kendall-Tau correlation. Heat maps illustrate the results for the various centrality measures under investigation. We use the Network package of Python for network construction and centralities calculation. We also compare the number of communities in each pair of cancerous and non-cancerous PPI networks for each tissue type. The results are represented in a histogram.

We also used the Louvain algorithm to uncover the communities and investigate the results’ stability. The results are similar. We identify the key genes at the network and community levels based on degree centrality. Genes with the highest average degree are identified at the community level. Based on the community size, the top five communities are considered at the community level. Five genes with the highest average degree are identified from each community. Those identified genes are then checked for their expression levels and compared their expression in non-diseased tissue. It is done using tnmplot [17].

**Fig 01.**
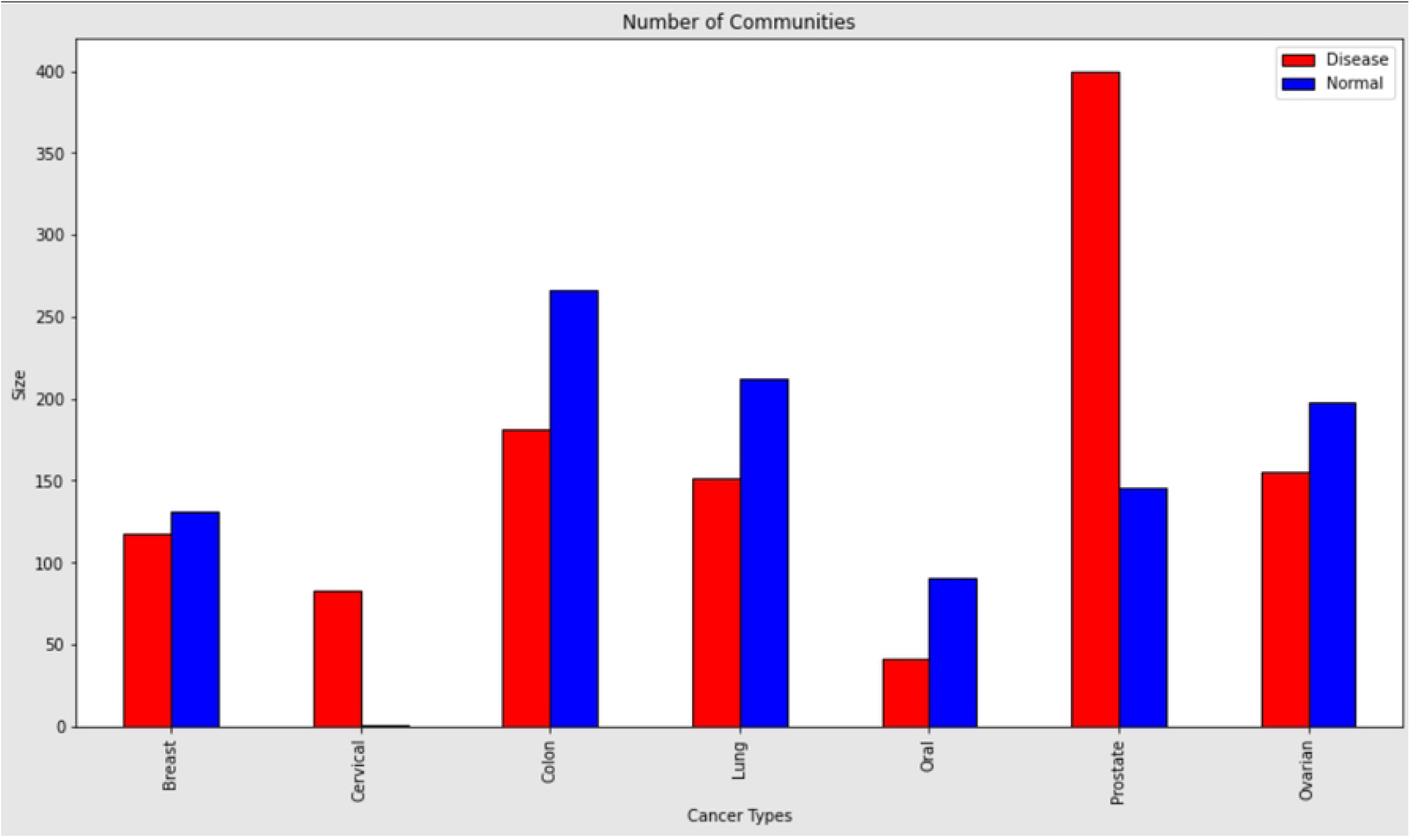
Histogram: The histogram compares the number of communities in the Cancerous and Non-Cancerous PPI networks

## Results

### Community Structure Analysis in the Cancerous and Non-Cancerous Datasets

Analyzing the mesoscopic structure, communities are generated using the Info map algorithm. In the majority of PPIs considered in the present study, the non-disease PPIs possess more communities than their respective cancerous PPIs except in cervical and prostate cancer. A maximum of four hundred communities are found in Prostate cancer PPI, whereas the maximum modularity value found is 0.7, in Prostate PPI. Among the non-cancerous PPIs, the highest number of communities found is two hundred and sixty-six, in the colon PPIs. The lung non-disease PPI holds the highest overall modularity score which is 0.74. The community structures are summarized in the following table 1 along with the modularity values. The comparison among the number of communities in the disease and non-diseased networks is represented through a histogram.

**Table 1.** This table shows the number of communities in different types of cancerous and non-cancerous tissues along with their modularity values.

| Cancer type | Number of Communities<br>in Disease data<br>(Modularity) | Number of Communities<br>in Non-Disease data<br>(Modularity) |
| --- | --- | --- |
| Breast Cancer | 118(0.56) | 131(0.54) |
| Cervical Cancer | 83(0.64) | 1(0.000) |
| Colon Cancer | 181(0.39) | 266(0.50) |
| Lung Cancer | 151(0.73) | 212(0.74) |
| Oral Cancer | 41(0.33) | 91(0.49) |
| Prostate Cancer | 400(0.674) | 146(0.70) |
| Ovarian Cancer | 155(0.62) | 198 (0.55) |

The Kendall-Tau correlation is computed among centrality measures like degree, betweenness, closeness, Katz, PageRank, and sub-graph at the network and community levels to compare the mesoscopic and macroscopic centralities. Community levels are further segregated into local and global, where the local considers the intra-community links and the global focuses on the inter-community links. The Kendall tau correlation is represented through heat maps for the cervical and prostate cancerous and non-cancerous PPIs. Among all the centrality measures computed, the classical degree centrality has consistently shown a good correlation with the local degree in all the PPI networks. The correlation between the classical and local degrees of prostate cancer PPI is 0.87 whereas, the correlation between classical and global modular degrees is found to be 0.6.

**Fig 02, Fig 03:**
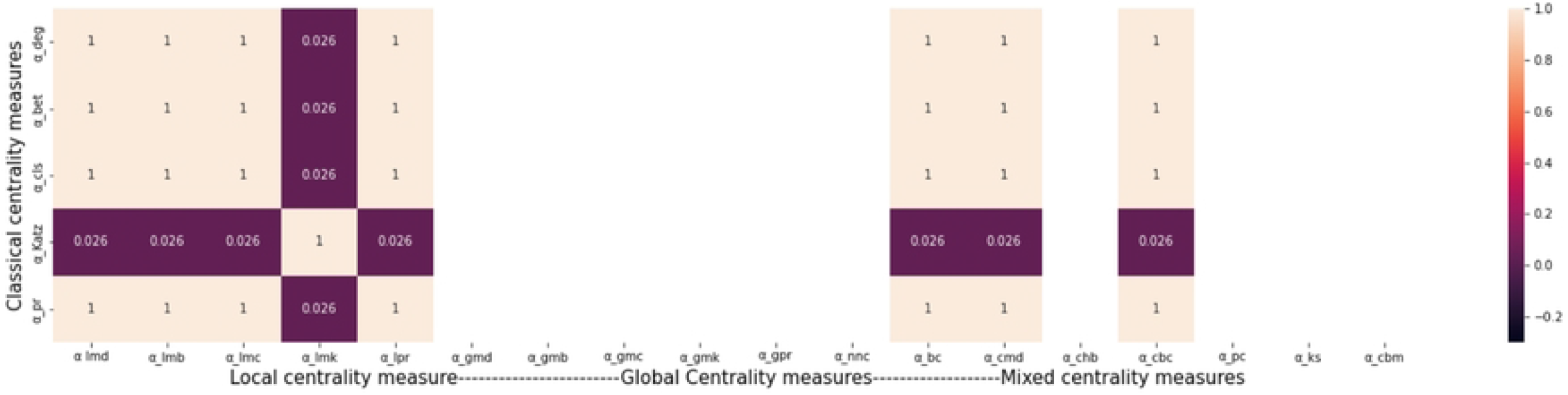
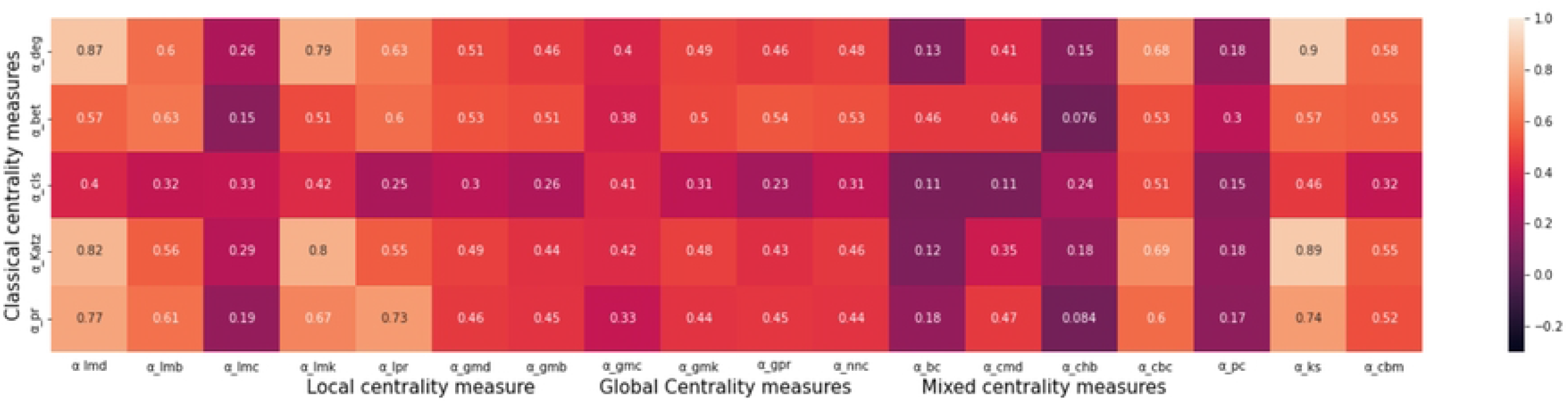
Heatmap: Kendall Tau’s Correlation heat map for Cervical Cancerous and Non-cancerous PPI networks. The top heat map shows the correlation between the classical centralities with the local and global centrality measures of the non-cancerous Cervical cancer tissue. The bottom represents the cancerous cervical Cells. The *α* deg refers to the classical degree, *α* bet is the classical betweenness, *α*ccls is classical closeness,*α* Katz classical Katz, *α* pr represents the classical PageRank. And the *α* lmd represents the local modular degree, *α* lmb represents the local modular betweenness,*α* lmc is the local modular closeness, *α* lmk is the local modular katz and *α* lpr is the local modular page rank and the global modular centralities are represented by *α* gmd global modular degree, *α* gmb global modular betweenness, *α* gmc is global modular closeness,*α* gmk global modular Katz, and *α* gpr represents the global modular page rank

**Fig 4, Fig 5:**
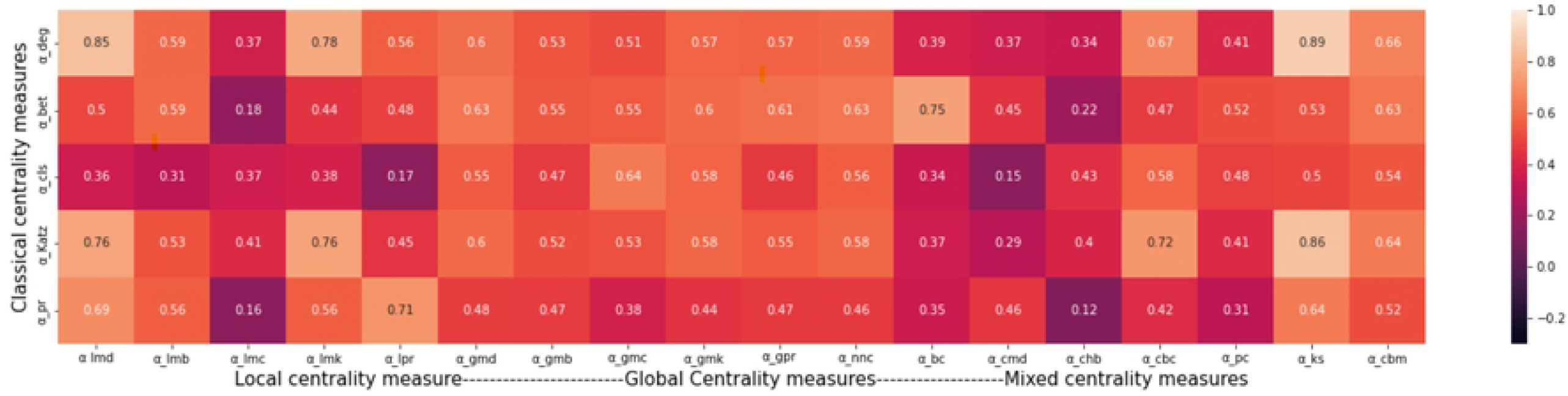
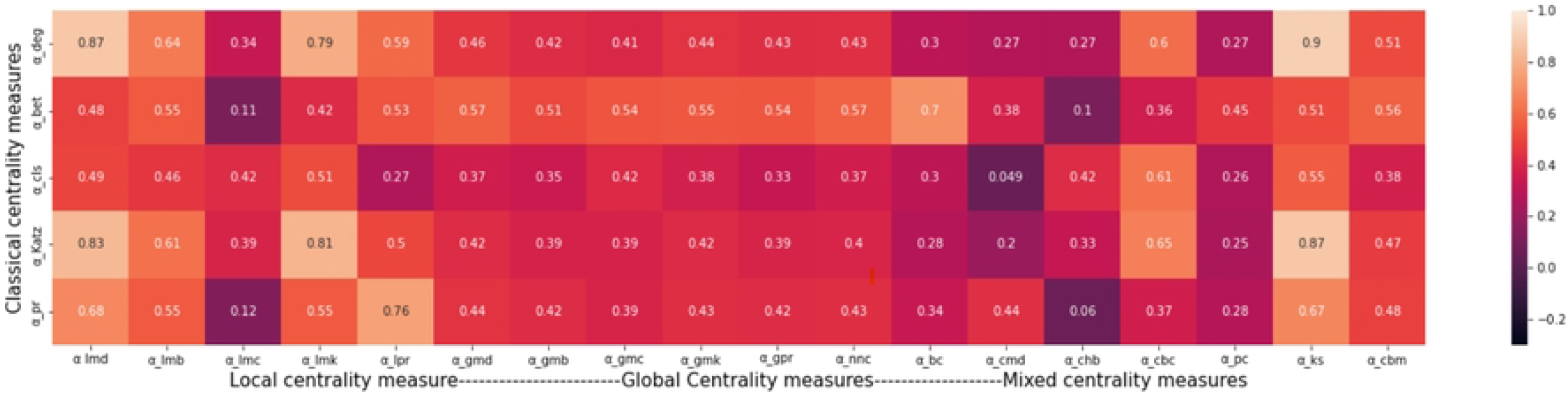
Kendall Tau’s Correlation heat map for Prostate Cancerous and Non-cancerous PPI networks. The above heat map shows the correlation of the non-cancerous Prostate cancer tissue and the bottom representation is for the Cancerous Prostate Cells.

To get an insight into the PPIs’ structural composition, the communities’ size in the diseased and non-diseased are compared. The results indicate that almost all the PPIs have few big-sized communities along with multiple small-sized communities, particularly in the Lung, ovarian, and oral cancerous tissues. The results are represented in the Figure The breast and colon cancerous and non-cancerous PPIs follow similar community structure patterns. A vast variation among the sizes of communities is observed, the largest community consists of 725 genes in the colon non-cancerous PPI, whereas multiple communities of minimum size two also exist in all the PPIs.

**Fig 06:**
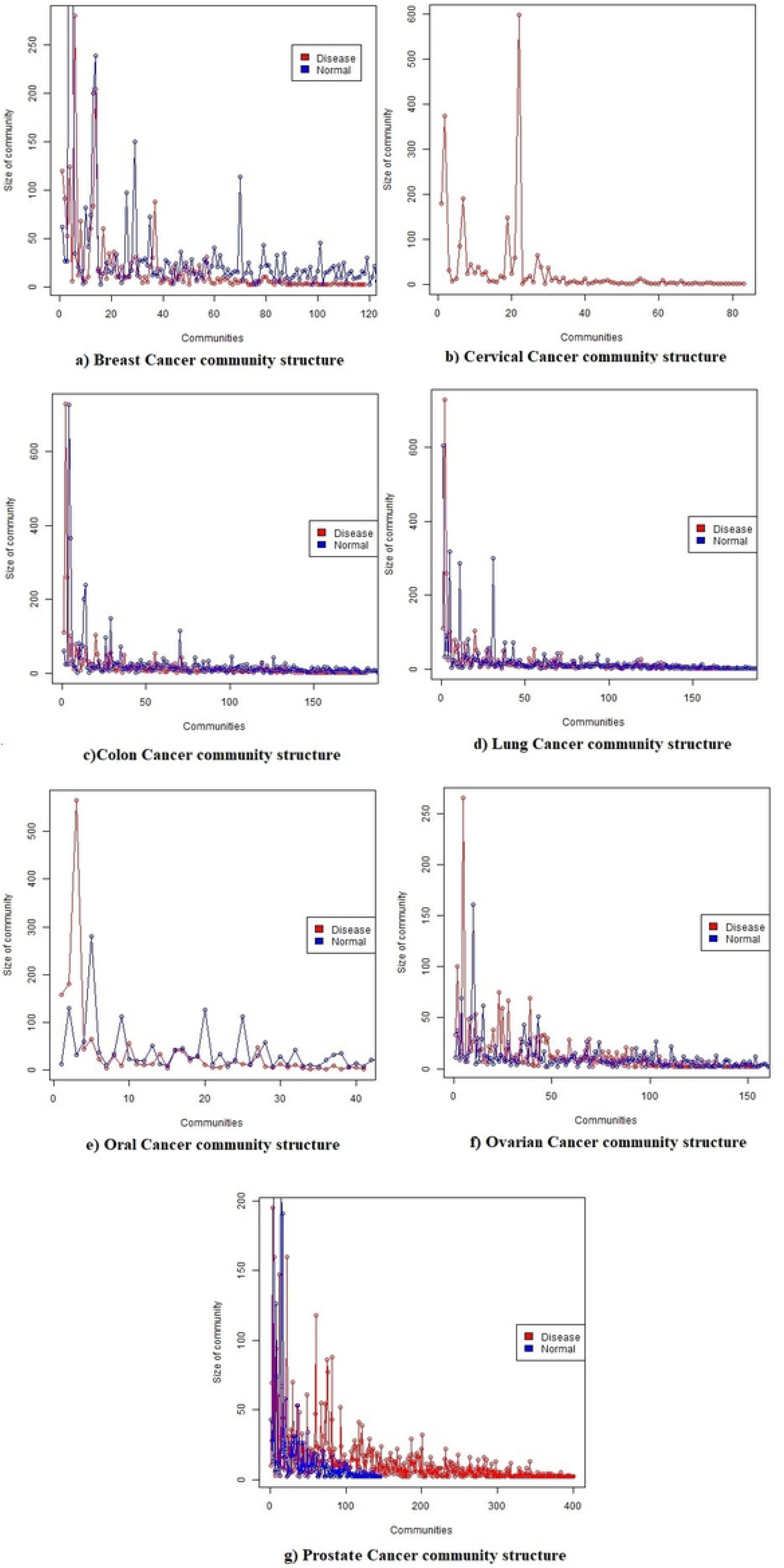
These graphs compare the sizes of communities in the different cancer types namely breast(a), colon(b), lung(c), prostate(d), oral(e), and ovarian(f). This shows the variation in the size of communities found in the different diseased and non-diseased PPIs

### Key gene Identification

Further analysis is directed to identify the key genes at the mesoscopic level. The results indicated that the PPIs of the non-diseased cells are more modular than their cancerous counterparts, except in the prostate and cervical, the rest of the investigation is conducted considering these two cancer types. The top five genes of the top five communities (w.r.t. the size of the community) are identified. Among all the top genes identified at the community level in cervical cancerous PPIs, the following genes are short-listed after observing the change in their genes expression level,**TP53, AKT1, BRCA1, SRC, PSMD14, MRPL3, NUP37, ESR1, and TRAPPC8**. The results indicate that the first five genes TP53, AKT1, BRCA1, SRC, PSMD14, MRPL3, and NUP37 are upregulated while the ESR1 and TRAPPC8 are downregulated. Among the identified genes TP53, AKT1, BRCA1, SRC, and ESR1 show a high degree-centric nature at the network level too, however PSMD14, MRPL3, NUP37, and TRAPPC8 are relatively silent at the network level. And they could prove to be potential drug targets after further research.

**Fig 7:**
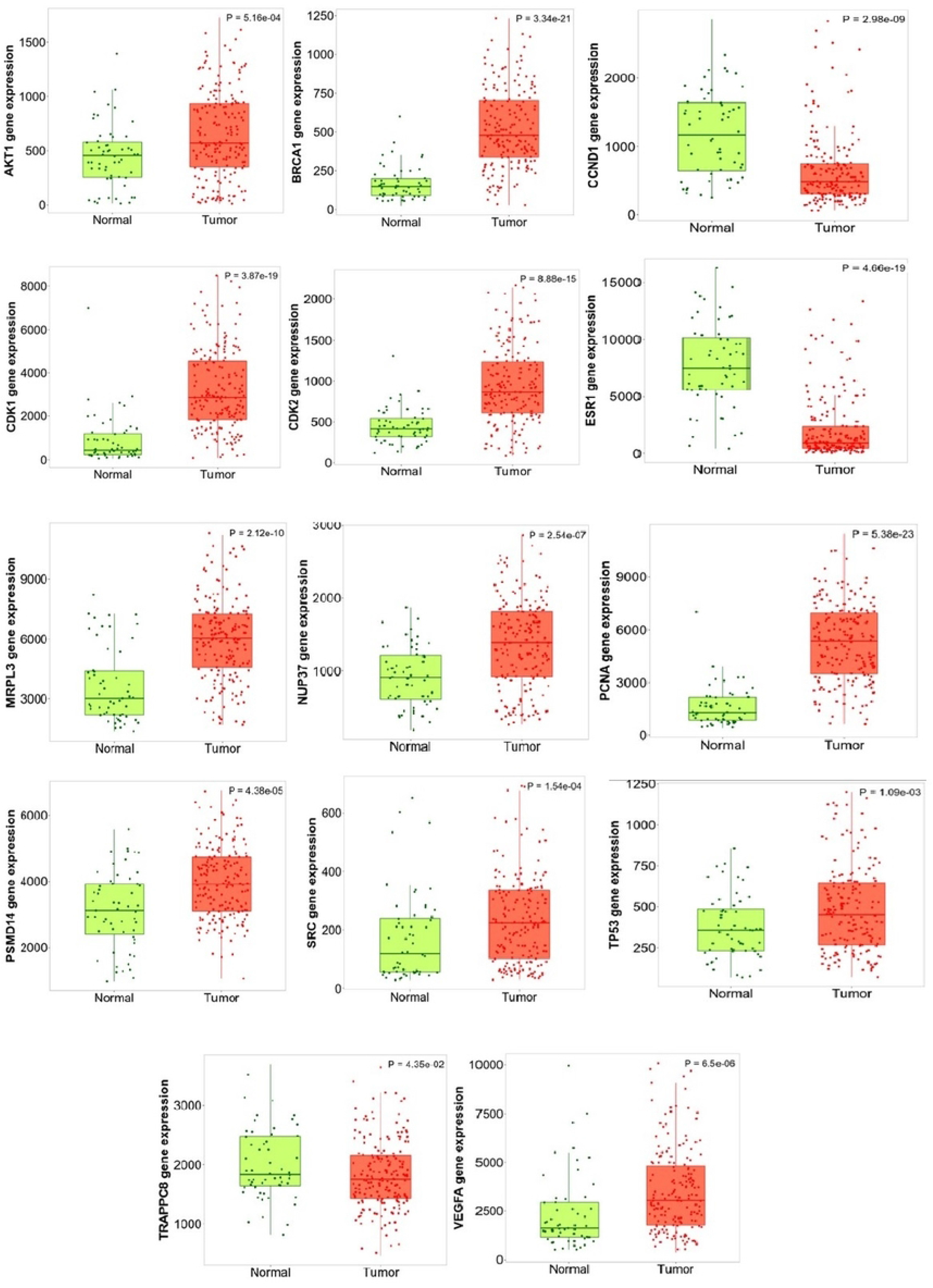
This figure shows the difference in the expression levels of the top identified genes in the cancerous and non-cancerous tissue of Cervical cancer. The box plot in green denotes the expression of the specified gene in normal tissue and the red box plot refers to the expression of the specified gene in the cancerous tissue. The box plots are generated using tnmplot [17]

Considering the prostate cancerous PPIs, the high-degree centric genes are identified at the community level and then screened for their expression levels. The following are the top genes found **FOS, TP53, UBA52, HLA-B, TSPO, CD19, INS, FYN, SUMO1, MDM2, and CCR5**. The expression level shows that **FOS, TP53, UBA52, HLA-B, TSPO, and CD19** are upregulated and **INS, FYN, SUMO1, MDM2, and CCR5** are down-regulated in the tumorous cell which can be seen in the figure Among these top genes TP53, UBA52, TSPO, and INS are found degree centric at the network level too, however, FOS, HLA-B, CD19, FYN, SUMO1, MDM2, and CCR5 show more interactivity at the community level.

**Fig 8:**
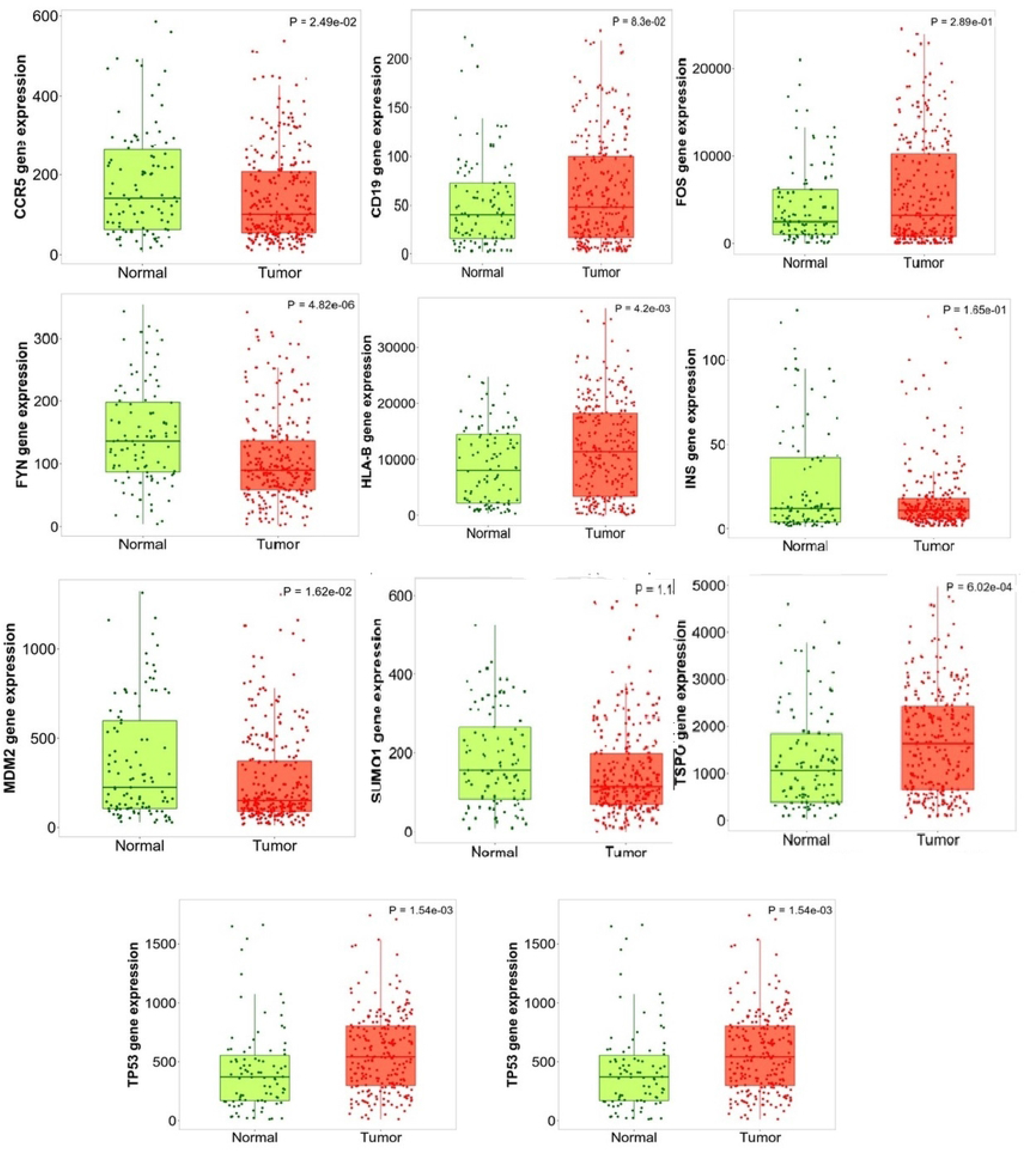
This figure shows the difference in the expression levels of the top identified genes in the cancerous and non-cancerous tissue of Prostate cancer. The box plot in green denotes the normal tissue and the red refers to the expression in cancerous tissue. The box plots are generated using tnmplot [17]

## Discussion

PPI network analysis has become a promising method to understand the mechanism of different diseases [18]. The PPI network analysis in the present study identifies the high degree centric genes at the community levels. Investigating modularity in biological networks is essential to understanding the interlinked nature of biological functional modules. Biological networks with high modularity refer to biological functions with fewer interactions. However, biological networks with less modularity refer to highly interlinked biological functions. The less connected networks tend to possess highly centric nodes that would be referred to as the network centrality measures. However, the networks with less connectivity among modules may tend to refer to strong community structures with fewer links among themselves. In the present situation, modularity plays a crucial role in understanding the importance of certain genes, within a given cancer network, among the two centrality settings i.e. mesoscopic and macroscopic. This study aimed to perform a comparative analysis among the two types of centrality measures in PPI cancer networks, the local and global community aware centrality measures. The correlation between the classical and local degree centrality is found high in almost all networks, so further investigation regarding the key gene identification is done considering the degree of centrality at the community level.

The modularity-based evaluation shows that PPI networks of non-diseased cells are more modular except for cervical and prostate cancer. The cervical and prostate cancer cells are found more modular, and the PPI network of Colon, Lung, Oral, Breast, and Ovarian cancer are found less modular than their respective non-cancerous PPI, in the studied dataset. This shows that cervical and prostate cancer possess likewise structural composition among the functional modules as compared to the other considered cancer networks. The prostate and cervical cancer networks are found with high correlations among the classical and local modular degree and betweenness centrality whereas the correlations among the closeness and global modular closeness are higher than the local closeness. As the results indicate that the correlation between, the classical degree and local modular degree and classical betweenness and local modular betweenness is high as compared to the global modular degree, it can be concluded that the network-based centralities cannot be the real representative centralities for the entire network in terms of a functional module. Instead, each functional module interpreted as a community possesses significantly different key genes in terms of the degree distribution, and edge betweenness in particular. Twenty-five genes were short-listed as hub genes, based on a high average degree at the community level for each cancerous PPI. These genes are further examined by checking their gene expression levels. Among the identified genes the following genes TP53, CDK1, AKT1, CDK2, BRCA1, VEGFA PCNA, SRC, PSMD14, MRPL3, and NUP37 are found to be upregulated in the cervical tumorous cells as ESR1, CCND1, and TRAPPC8 are downregulated, which confirms their behavioral change in the diseased and non-diseased network. The gene expression level is examined using the online tool tnmplot [17].

Several approaches have been adopted to identify key genes in different cancers including the comparison of disease and non-diseased networks [19] and BI analysis etc. The following genes TSPO, CCND1, and FOS were found to be down-regulated while CDK1, TOP2A, CCNB1, PCNA, BIRC5, and MAD2L1 up-regulated in cervical cancer through a BI analysis [20]. Similarly, these twelve genes were identified as cervical cancer key gene ECT2, WDHD1, TYMS, FANCI, CXCL8, MCM2, HELLS, DTL, CEP55, CDK1, AURKA, TOP2A using the BI analysis [21]. While exploring the prognosis of cervical cancer BTNL8, CCR7, CD1E, CD6, CD27, CD79A, GRAP2, SP1B, and LY9 were identified as the bio-markers for the cervical cancer diagnosis, prognosis, and therapeutics as well [22]. The association between the genes and the clinical characteristics of prostate cancer was studied using the weighted graph correlation network analysis (WGCNA) and the following hub genes LMNB1, TK1, RACGAP1, and ZWIN were identified [23]. Through BI analysis on PPI, [24] found RPS21, FOXO1, BIRC5, POLR2H, RPL22L1, and NPM1 as the prostate cancer key genes. CDK1, CCND1, PCNA [20], CDK2, AND VEGFA [25], PLOD2, DSG2, SPP1, CXCL8, MCM5, HLTF, and KLF4 [26], KNTC1 [27], CDKN2A, IL1R2 and RFC4 [28], and AKT1 ‘ [29] have been reported to have some significance in cause or prognosis of cervical cancer. The different roles of CDK1, CCNB1, ITGB1, FN1, MMP9 and STAT1 are also being discussed in the literature [30].

According to our analysis TP53, BRCA1, SRC, PSMD14, MRPL3, NUP37, ESR1, and TRAPPC8 are top genes in cervical cancer. TRAPPC8 is a part of the trafficking protein particle complex and is involved in Golgi organization [31]. ESR1 encodes an estrogen receptor and ligand activation. ESR1 encoded receptors are known to play an important role in breast cancer, endometrial cancer, and osteoporosis [31], loss of ESR1 is reported to increase the cervical cancer occurrence [32]. NUP37 is a nuclear pore complex responsible for macromolecule transportation between the nucleus and cytoplasm [31]. MRPL3 mitochondrial ribosomal proteins are nuclear-encoded genes. They assist in protein synthesis within mitochondrion [31]. PSMD14 encodes a component of 26S proteasome, a multiprotein complex. The degradation of ubiquitinated intracellular proteins is catalyzed by it. SRC is a proto-oncogene that may regulate the embryonic development of and cell growth [31]. The SRC-ERK signaling pathway may prove to be a favorable therapeutic target against cervical cancer [33]. BRAC1 encodes a nuclear phosphoprotein which plays its part in maintaining genomic stability. It is also known as a tumor suppressor. Mutation in this gene is associated with 40 percent of inherited breast cancer [31]. Though BRCA1 is mostly known for its association with breast cancer, it is studied as having its role in cervical cancer as well [33]. TP53 is known to be mutated in most cancers [31]and in our results also been found in more than 50 percent of cancer types that we explored during our experimentation.

The top genes identified at the mesoscopic level in prostate cancer cells when checked for the expression level found that FOS, TP53, UBA52, HLA-B, TSPO, and CD19 are upregulated in the tumorous cells whereas INS, FYN, SUMO1, and CCR5 are down-regulated in a diseased cell. The change in expression levels of the identified top genes validates our finding. The results are shown in Figure 3. FOS is a proto-oncogene which implies that mutation in it can cause cancer. These proteins are involved in cell transformation, differentiation, and proliferation [31]. The network analysis performed on PPI of prostate cancer and normal samples revealed UBA52 and SUMO2 are identified as key genes [34]. The increased expression of TSPO has been reported earlier, and its correlation with prostate cancer progression. CD19 and INS have been reported as hub genes while inspecting at the network level [35], and these two are identified as our hub genes at the community level as well. FYN shows high connectivity at the community level in our results, and it has been reported to have high expression levels in prostate cancer [35]. SUMO1 encodes a protein of the SUMO(small ubiquitin-like modifier) family, which is involved in various cellular processes like transcriptional regulation, nuclear transport protein stability, and apoptosis [31]. CCR5 is found on the white blood cell’s surface and is involved in the immune system, a chemokine receptor. The top genes found in this research work are the key genes identified for different cancers that validate our work and more research in this direction can be useful in identifying new drug targets through the identification of key genes.

## Conclusion

Network analysis is applied to the protein-protein interaction(PPI) network to get insight on the structural composition of the biological network at the macro and mesoscopic levels. Our findings indicate that a lot can be explored by studying the biological networks at the community level. The results indicate that the diseased and non-diseased PPI networks are highly modular. The correlation among different centrality measures indicates that the important genes at the network and community levels might differ. The genes may play an active role at the community level while being silent at the network level. Certain biological processes may have distinct unique key genes at the community level and that key gene can’t be ignored in therapeutics. It may be concluded that the gene’s significance at the mesoscopic level may guide us more toward potential drug targets. In this dataset, prostate and cervical cancerous cells’ PPIs are found to be more modular than their respective normal cells’ PPIs. The hub genes at the community level are identified for these two cancerous networks.**TP53**,**BRCA1, SRC**, **PSMD14**, **MRPL3**, **NUP37**, **ESR1 and TRAPPC8** are found as top genes in cervical cancer whereas **FOS**, **TP53**, **UBA52**, **HLA-B**, **TSPO and CD19**, **INS, FYN, SUMO1, and CCR5** are found as top genes in prostate cancer. As the results show that the normal tissue PPIs are more modular as compared to the cancerous tissue PPI in most networks except prostate and cervical, then it could be interpreted that certain genes were not active or regulated in the normal condition and now they become active and bridged the communities.

